# Polyethylene terephthalate (PET) micro- and nanoplastic particles affect the mitochondrial efficiency of human brain vascular pericytes without inducing oxidative stress

**DOI:** 10.1101/2023.10.24.563735

**Authors:** Sean M. Gettings, William Timbury, Anna Dmochowska, Riddhi Sharma, Lewis E. MacKenzie, Guillaume Miquelard-Garnier, Nora Bourbia

## Abstract

The objective of this investigation was to evaluate the influence of micro- and nanoplastic particles composed of polyethylene terephthalate (PET), a significant contributor to plastic pollution, on human brain vascular pericytes. Specifically, we delved into their impact on mitochondrial functionality, oxidative stress, and the expression of genes associated with oxidative stress and ferroptosis. Our findings demonstrate that the exposure of a monoculture of human brain vascular pericytes to PET particles *in vitro* at a concentration of 50 ppm for a duration of 6 days did not elicit oxidative stress. Notably, we observed an augmentation in various aspects of mitochondrial respiration, including extracellular acidification, proton pump leakage, maximal respiration, spare respiratory capacity, and ATP production in pericytes subjected to PET particles. Furthermore, there were no statistically significant alterations in mitochondrial DNA copy number, or the expression of genes linked to oxidative stress and ferroptosis.

These outcomes suggest that, at a concentration of 50 parts per million (ppm) and for 6 days exposure, PET particles do not induce oxidative stress in human brain vascular pericytes. Instead, they seem to incite a potential mitochondrial hormesis, also named mitohormesis, response, which seemingly enhances mitochondrial function. Further investigations are warranted to explore the stages of mitohormesis and the potential consequences of plastics on the integrity of the blood-brain barrier and intercellular interactions. This research contributes to our comprehension of the potential repercussions of nanoplastic pollution on human health and underscores the imperative need for ongoing examinations into the exposure to plastic particles.

**Graphical Abstract (created with BioRender.com):** 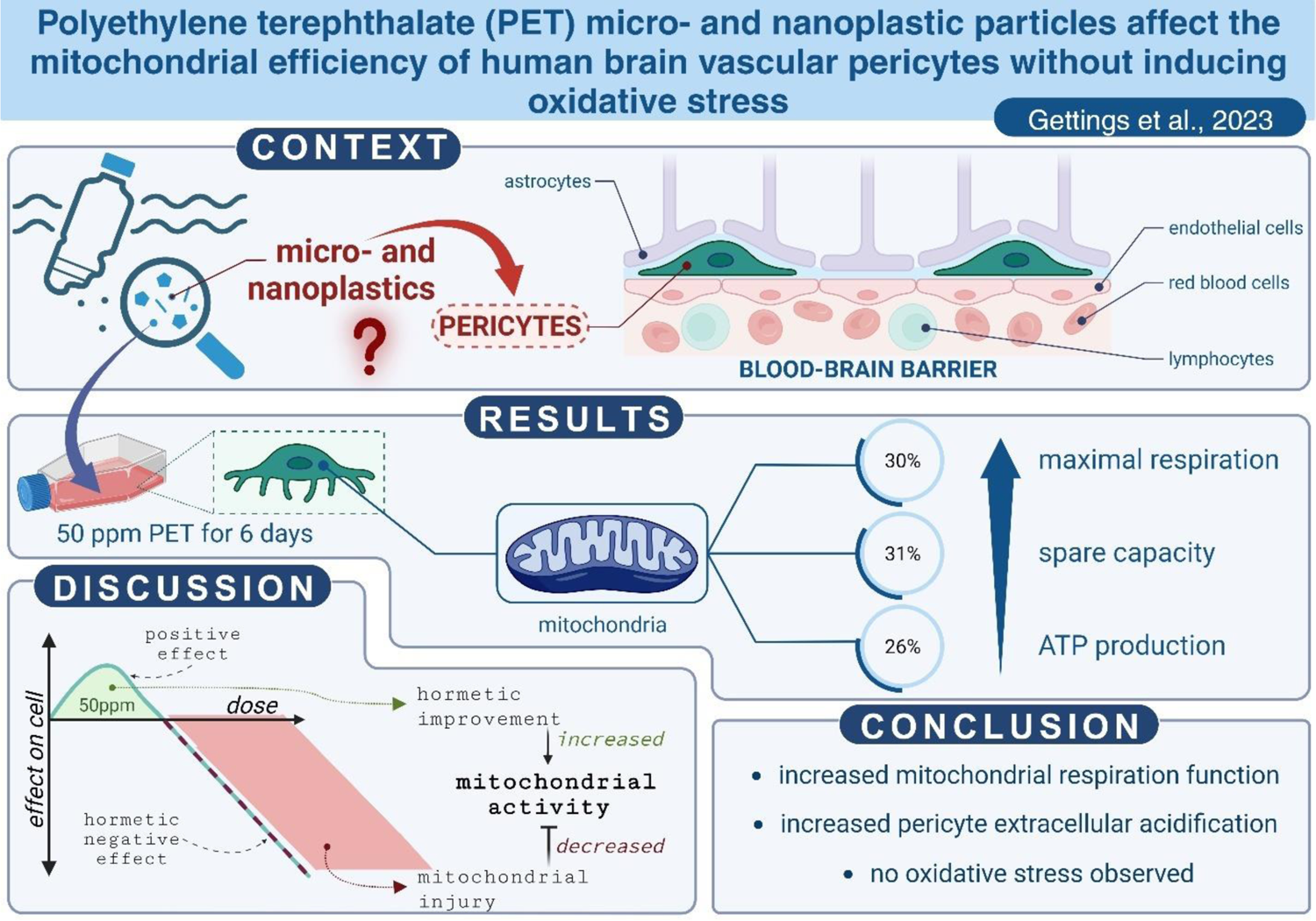

**Highlight:**

- Fabrication of polyethylene terephthalate (PET) micro- and nanoplastics
- PET particles increase pericytes mitochondrial respiration functions
- PET particles increase pericytes extracellular acidification
- Oxidative stress was not observed in pericytes subjected to PET particles

## 1. Introduction

Nanoplastics (< 0.1 µm) and microplastics (< 5000 µm) are posing a health threat as they are polluting our environment (Landrigan et al., 2020; Zhang et al., 2021) and food (Liu et al., 2021) to the point of finding plastic particles in human blood (Leslie et al., 2022), lung (Jenner et al., 2022), placenta (Ragusa et al., 2021), heart (Yang et al., 2023), and breast milk (Ragusa et al., 2022). Plastic particles are also released from plastic packaging (Jadhav et al., 2021). For instance, it has been estimated that microwaving plastic containers for three minutes could release about 4 million microplastics and about 2 billion nanoplastic particles from one square meter of plastic (Hussain et al., 2023). The same authors estimated the highest daily intake of plastic particles from microwaved water or dairy products in plastic containers alone represents about 20 ng/kg per day for toddlers consuming microwaved dairy products from polypropylene containers.

Studies on the effect of nanoplastics on brain and blood-brain barrier functions are still limited (Prüst et al., 2020) but it is established that plastic particles cross the blood-brain barrier and can be transferred through the food chain (Kopatz et al., 2023; Mattsson et al., 2017). Additionally, nanoplastics can be carriers for other molecules to enhance their accumulation inside the brain (Chen et al., 2017). Plastic particles can also induce oxidative stress (Lei et al., 2018; Schirinzi et al., 2017), mitochondrial dysfunctions (Murali et al., 2015), and affect the expression of genes linked to neurodevelopment in a 3-dimensional model of human embryonic stem cells. This included genes associated with neurogenesis regulation and telencephalon differentiation (Hoelting et al., 2013).

Pericyte cells are fundamental constituent of the blood-brain barrier and have several critical functions such as maintaining the blood-brain barrier and clearing toxins (Bell et al., 2010; Quaegebeur et al., 2010). Their dysfunctions are associated with neurodegenerative diseases such as Alzheimer’s and Parkinson’s diseases (Baloyannis and Baloyannis, 2012; Halliday et al., 2016; Lendahl et al., 2019; Quaegebeur et al., 2010; Winkler et al., 2014), which are multifactorial diseases with where environmental pollutants (e.g. neurotoxic metals, metal-nanoparticles, and pesticides) are implicated as known risk factors (Chin-Chan et al., 2015; Kriit et al., 2021). Therefore, it is important to assess whether nano and microplastic particles can interfere pericytes, hindering their neuroprotective functions, particularly through the carrier molecule capacity of nanoparticles.

To experimentally study potential adverse effects of plastic particle exposure on pericytes, we made our own polyethylene terephthalate (PET) plastic particles. PET was selected as the plastic feed stock because it is widely used for food containers and drink bottles. Also, containers made of PET release more plastic than those made of polypropylene (Hussain et al., 2023). In study design, we confirmed the composition of PET particles using Fourier-transform infrared spectroscopy (FTIR), thermogravimetric analysis (TGA), and differential scanning calorimetry (DSC). Additionally, the concentration of plastic micro/nano particles is a key variable. We used a concentration of 50 parts per million of PET particles, i.e. 50 µg/ml which is 31.25 times higher than the average plastic particle detected in human blood (Leslie et al., 2022) to take in account potential bioaccumulation at the BBB (Shih et al., 2018) and occupational exposures (Salthammer, 2022).

Finally, we assessed whether PET particles can induce oxidative stress, mitochondrial dysfunctions, and eventually ferroptosis cell death on human brain vascular pericytes (HBVP) cultured *in vitro*. We aimed to simulate chronic exposure by exposing the cells to PET particles for 6 days.

### 2. Materiel and Methods

### 2.1. Plastic particle characterisation

#### 2.1.1. Plastic particle generation

Realistic plastic particles were generated based on Ji et al., 2020 with the adaptations as follows. Micro- and nanoplastic particles were generated from polyethylene terephthalate (PET) confectionary-style plastic jars purchased from a scientific supply retailer (cat: 129-0590; VWR International). These jars were selected as they can be readily purchased by other researchers to aid in comparative studies. The jars were then shredded using a hole-puncher to generate feed material (∼4 mm diameter). The PET feed material was ground in a Pulverisette 2 automatic pestle from the manufacturer Fritsch for 4 hours. The automatic pestle was cooled with dry ice (which was added every 15 minutes). The PET plastic was then placed in 7 ml steel tubes with 10 x 2 mm steel ball bearings to be broken down further with the Precellys Evolution Touch tissue homogeniser with the Cryolys Evolution attachment. The tubes containing PET were then purged with liquid nitrogen for 12 minutes before agitation in the Cyrolys Precellys at two cycles of 10,000 RPM for 60 seconds with a 20 second rest between cycles. This was repeated 5 times. The resulting PET micro- and nanoplastic mix was sieved from the remaining feed material using Wstyler brass sieves (No.60) down to a 250 µm tolerance. A stock concentration of 10,000 ppm PET with a particle size range of 82 nm to 250 µm was added to 0.05 % bovine serum albumin (BSA) (cat: A9647-10G; Merck) to facilitate the suspension and dispersal of PET particles.

#### 2.1.2. Plastic particle sizing

For scanning transmission electron microscopy (STEM) and scanning electron microscopy (SEM) imaging, a fresh grid (holey carbon film on 200 mesh, AGS147H, Agar Scientific Ltd) was gripped using tweezers and immersed within a freshly agitated particle solution. The grid was then air dried and then mounted on a copper grid holder and affixed with quick drying conductive silver paint (AGG302, Agar Scientific Ltd). Imaging was conducted using a variable pressure field emission gun scanning electron microscope (FEI Quanta 250) operating in high vacuum mode with an accelerating voltage of 30 kV and at 100,000X magnification. STEM imaging was conducted via a two-segment solid-state detector holder operating in bright-field mode and mounted under the sample. SEM imaging was conducted using the standard secondary electron (SE) detector. Images were saved in uncompressed .TIF format at a resolution of 1024 **×** 943 pixels. Nanoparticle diameters were measured manually from SEM images by using ImageJ software (Schneider et al., 2012).

#### 2.1.3. Plastic particle characterisation

The chemical composition of PET particles was evaluated by Fourier-transform infrared spectroscopy (FTIR). The sample was ground, mixed with dried spectroscopic grade potassium bromide, and formed into a pellet using a laboratory press. The mass of PET particles was equal to 1 weight percent (wt%) of the pellet. The spectrum was recorded on a Frontier spectrometer (Perkin Elmer) in the range of 4000-650 cm^-1^ at 2 cm^-1^ resolution and is an average of 32 scans.

Thermal stability of the sample was determined by thermogravimetric analysis (TGA) in the range of 25-600°C with a 10 deg/min heating rate (Q50, TA Instruments). The experiment was conducted under nitrogen atmosphere (50 mL/min) in a platinum crucible with a sample mass of about 7 mg.

Thermal phase transitions of the PET particles were analyzed by differential scanning calorimetry (DSC) in the range of 30–325 °C at 10 deg/min heating rate (Q1000, TA Instruments). Two heating cycles were applied in order to erase the thermomechanical history of the material. The measurement was performed under nitrogen flow (50mL/min) in a sealed aluminum pan with the sample mass of about 3 mg.

### 2.2. Cell culture

Human Brain Vascular Pericytes (HBVP) (cat.: 1200; ScienCell) were cultured in Pericyte Medium (PM, Cat. #1201-b; ScienCell) supplemented with 2 % foetal bovine serum (FBS) and 1 % Penicillin-Streptomycin in T25 flasks. At confluency 60-70 %, pericyte cells were exposed to either a final concentration of 50 ppm PET (average particle size: 205 nm) or the control medium (supplemented pericyte medium mentioned above) used for PET dilution for 6 days. At 6 days cells were processed for further experiment. Cells used for protein, DNA or RNA extractions were rinsed with Dulbecco’s phosphate-buffered saline (DPBS) (cat: 14040174, ThermoFisher), trypsinised (cat: T4174–100 mL, Merk) and the trypsin was neutralised using Pericyte medium supplemented as mentioned above. The cells were then centrifuged at 1000g, and the cell pellets collected for further experiments.

### 2.3. Cell counting

After 6 days treatment, cells were counted as described previously (Sharma et al., 2023) by washing the cells with DPBS, tripsynised and neutralised with pericyte medium supplemented with FBS. The cells were then counted using a C-Chip haemocytometer (cat: DHCN01, Scientific Laboratory Supplies).

### 2.4. Mitochondrial functions

#### 2.4.1. Seahorse XF Cell Mito Stress Test

To assess the mitochondrial respiration, a mitochondrial stress test was performed using the Agilent Seahorse XFp analyzer. The manufacturer protocol was followed using Seahorse XF Cell Mito Stress Test Kit (cat.: 103010-100; Agilent), Seahorse XFp FluxPak (103022-100; Agilent), and the Seahorse XF DMEM assay medium pack (103680-100; Agilent). Cell density of 40,000 cells per well, and drug concentrations (Oligomycine at 1.5 µM, Carbonyl cyanide-4 (trifluoromethoxy) phenylhydrazone at 0.5 µM, and rotenone/antimycin A at 0.5 µM) were determined and optimised for the HBVP cells prior to the experiment. Oxygen consumption rate (OCR) expressed in pmol/min is used to assess each aspect output of the Seahorse XF Cell Mito Stress except the coupling efficiency also expressed as a percentage, and the extracellular acidification (ECAR) is expressed in mpH/min.

#### 2.4.2. Relative mitochondrial DNA copy

To determine relative mitochondrial DNA copies of pericytes after exposure to plastic particles, DNA was extracted by applying DNeasy Blood & Tissue Kit (cat.: 69504; Qiagen) to the pelleted cells, followed by quantitative polymerase chain reaction (qPCR) using PerfeCTa SYBR Green SuperMix (cat.: 733-1246; VWR International) for an initial 95 ⁰C for 10 minutes followed by 40 cycles (95 ⁰C for 15 sec, followed by 62 ⁰C for 60 sec, and then 72 ⁰C for 20 sec) in a Magnetic Induction Cycler (Mic qPCR,Bio Molecular Systems).

The primer set of the nuclear DNA was selected to target the gene *B2M* (alias Beta-2-Microglobulin) (Forward sequence: 5’ TGCTGTCTCCATGTTTGATGTATCT 3’; Reverse sequence: 5’ TCTCTGCTCCCCACCTCTAAGT 3’; the amplicon size is 86 base pairs) and the primer set for the mitochondrial DNA was selected to target the gene *MT-TL1* (alias TRNA-Leu (UUR)) (Forward sequence: 5’ CACCCAAGAACAGGGTTTGT 3’; Reverse sequence: 5’ TGGCCATGGGTATGTTGTTA 3’; the amplicon size is 107 base pairs). Both set of primers have been validated for their efficiency prior experiment by testing a standard curve of each primer pairs using a 1:3 serial dilution of pericyte DNA. The relative mitochondrial DNA content was determined as the formula 2 x 2^ΔCT^, where ΔCT = nuclear DNA CT - mitochondrial DNA CT (Rooney et al., 2015).

### 2.5. Oxidative stress responses and ferroptosis

#### 2.5.1. Oxidative stress assay

The presence of reactive oxygen species (ROS) was assessed using the general oxidative stress indicator CM-H2DCFDA (cat: C6827, ThermoFisher Scientific). As per the manufacturer’s protocol, the CM-H2DCFDA probe was initially reconstituted in dimethyl sulfoxide (DMSO) (cat: 317275, Sigma-Aldrich) and then further dilutions were carried out using PM to achieve a final concentration of 5 µM. Cells were washed using 1 ml HBSS (cat: H6648, Sigma-Aldrich) per T25 flask, then given 5 ml of the probe for a 45 minute – 1 hour incubation period. After which, the cells were trypsinised with 1 ml Trypsin and subsequently quenched with 2 ml DTI (cat: R-007-100, ThermoFisher Scientific). Cells were then placed into falcon tubes and centrifuged at 1000 g at 4 °C for 5 minutes before the supernatant was aspirated and the cell pellet was resuspended in 1 ml DPBS. From this 100 µl of the cell suspension was seeded in sextuplicate on a 96 well-plate along with 150 µl of propidium iodide at a concentration of 50 µg/ml. Positive controls bookended each row and were achieved by adding hydrogen peroxide (cat: 101284N, VWR International) to achieve a final concentration of 0.3 %. Samples were then allowed to incubate for a further 15 minutes before running the assay. A total of 5000 events per wells were analysed with a Guava EasyCyte HT flow cytometer (Merk Millipore) from which cells positive for propidium iodide were excluded from further analysis. The geometric mean fluorescence intensity was compared between the experimental groups.

#### 2.5.2. Expression of genes associated to oxidative stress and ferroptosis

Reverse transcription quantitative polymerase chain reaction (RT-qPCR) was used to assess whether the PET particles had an impact on the expression of the genes associated or impaired during ferroptosis and/or oxidative stress. RNA extraction of the pelleted cells was done using RiboPure™ RNA Purification Kit (cat.: AM1924; Thermo Fisher Scientific) followed by a cDNA conversion using High-Capacity cDNA Reverse Transcription kit (cat.: 4368814; Thermo Fisher Scientific). RT-qPCR was performed using PerfeCTa SYBR Green SuperMix with the equipment Mic qPCR for initial 90 ⁰C for 2 min followed by 40 cycles (95 ⁰C for 5 sec, followed by 58 ⁰C for 15 sec, and then 72 ⁰C for 10 sec). All set of primers (Table 1) were previously validated for their efficiency in prior experiments by testing a standard curve of each primer pairs using a 1:3 serial dilution of pericyte cDNA. The gene beta-2-microglobulin (*B2M*) is used as the housekeeping gene to determine the delta cycle threshold (ΔCT) which was then used to perform statistical analysis to compare both experimental groups.

**Table 1.**
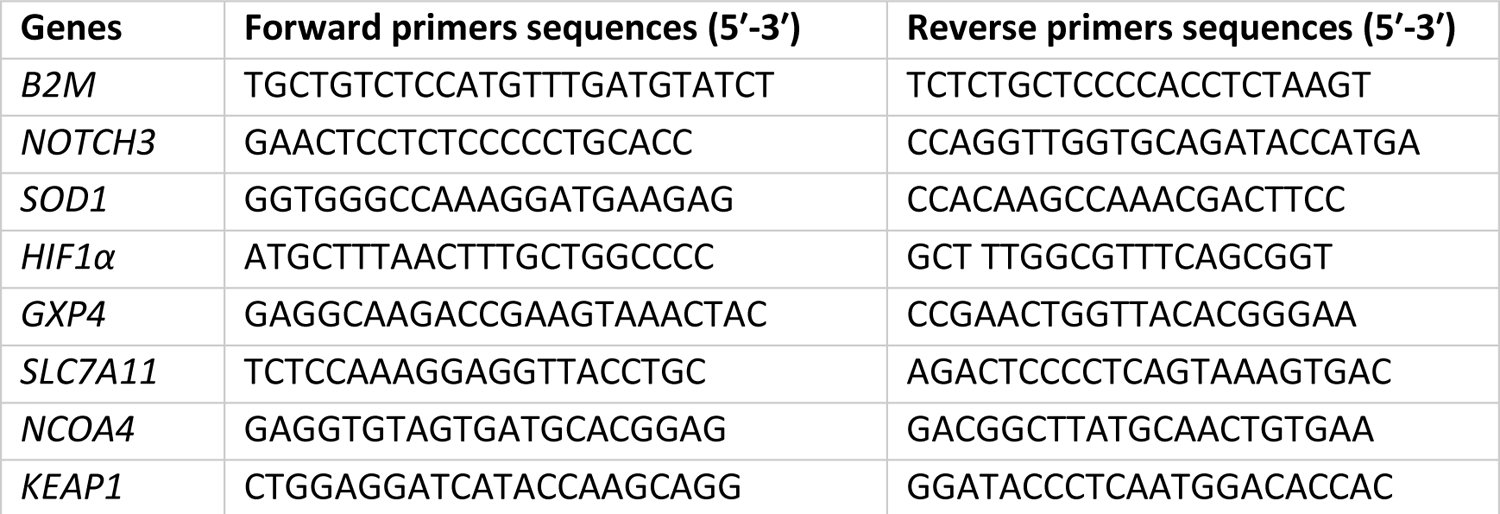
List of rt-qPCR primer sequences.

### 2.6. Statistics

Normality distribution between samples was tested using the Shapiro-Wilk test. The Mann-Whitney U test was used for non-parametric comparison between experimental groups, and the two-tailed unpaired t-test was used when comparing experimental groups with a normal distribution. 2-way ANOVA was used for multiple comparisons between both experimental groups. All statistical analysis has been performed using the software GraphPad Prism 9.4.0 and all figures were formatted in Inkscape V.1.1.2.

## 3. Results

### 3.1. Plastics characterisation

From electron microscopy (Figure 1), the average diameter of the PET particles observed was determined to be 205 ± 79 nm (diameter ± std. deviation; n = 60) with a particle size ranging from 82 nm to 385 nm. This confirmed that the 10000 ppm PET stock contains nanoparticles (nanoparticles are defined as particles < 1000 nm) (Hartmann et al., 2019). Microplastics were observed in the medium however it was apparent they did not adhere to the electron macroscopy grid as no microplastic particles were observed under electron microscopy.

**Figure 1.**
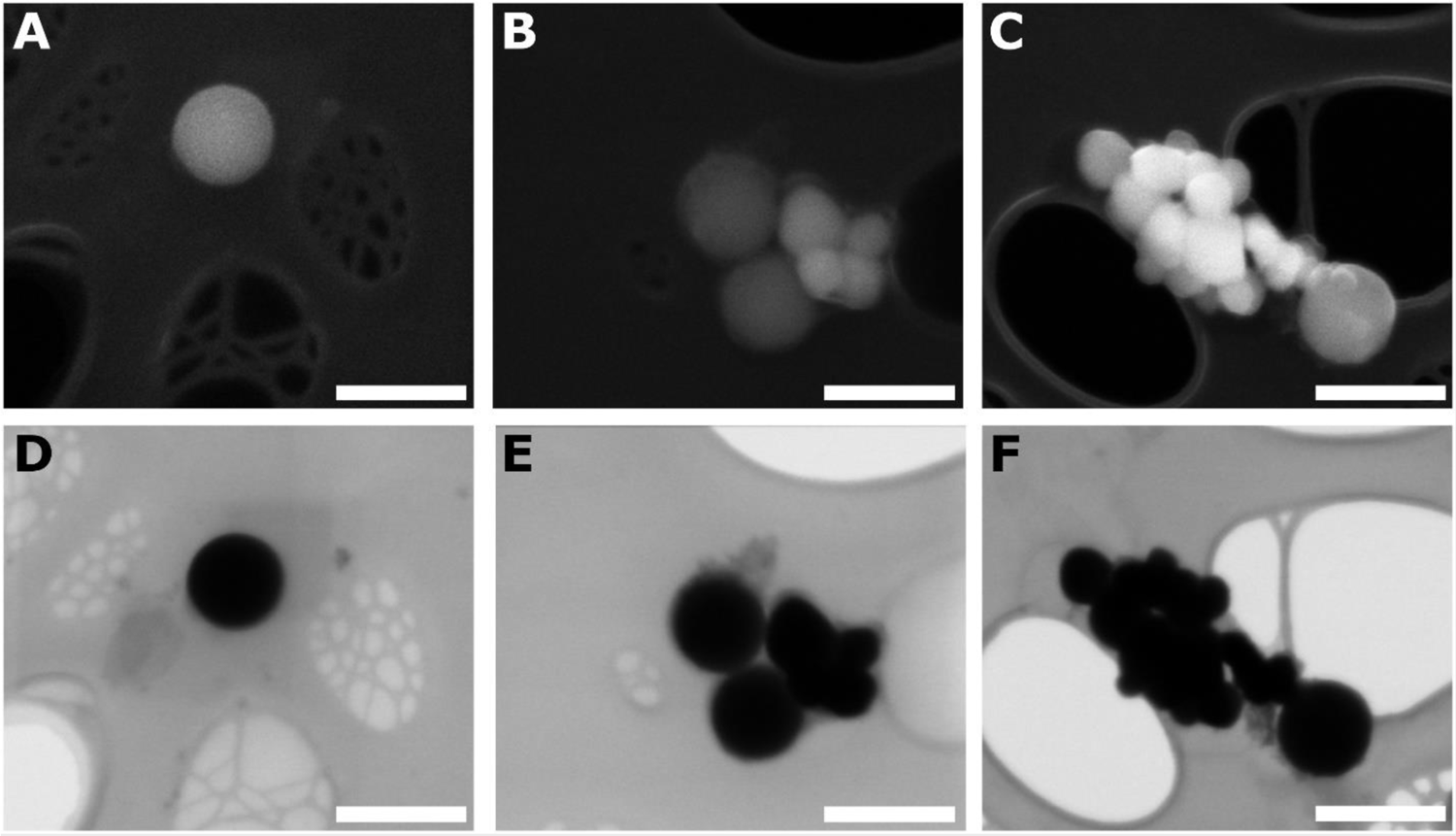
Representative SEM (top row; A, B and C) and STEM (bottom row; D, E and F) images of nano plastics used in this study. Scale bar represents 400 nm. SEM more clearly shows the morphology of individual and agglomerated nanoplastics.

The FTIR results clearly confirm that PET particles have undetectable level of impurities as only peaks correlated with the structure of PET were distinguished in the spectrum (Figure 2A). The peak at 2970 cm^-1^ was attributed to C–H stretching vibrations (Fávaro et al., 2007). Carbonyl stretching band can be seen at 1717 cm^-1^, corresponding to C=O stretching. Skeletal stretching vibrations were observed at 1410 cm^-1^. The peak at 1344 cm^-1^ corresponded to –CH_2_ wagging in trans conformation (Andanson and Kazarian, 2008). Three different vibrations can be seen at 1244 cm^-1^, namely the stretching of C(–O)O, stretching of ring-ester C–C, and in-plane bending of C–O (Donelli et al., 2009). The peaks at 1122 and 1098 cm^-1^ were correlated with C–O stretching. This doublet is linked with the crystallinity of PET. The higher the intensity of the peak at 1122 cm^-1^, the bigger the amount of the crystalline phase. The band at 1011 cm^-1^ indicated the in-plane vibrations of benzene. Finally, the out-of-plane benzene group vibrations were observed at 871 and 723 cm^-1^. This result confirms that no significant amount of additives are present in the obtained nanoparticles.

**Figure 2.**
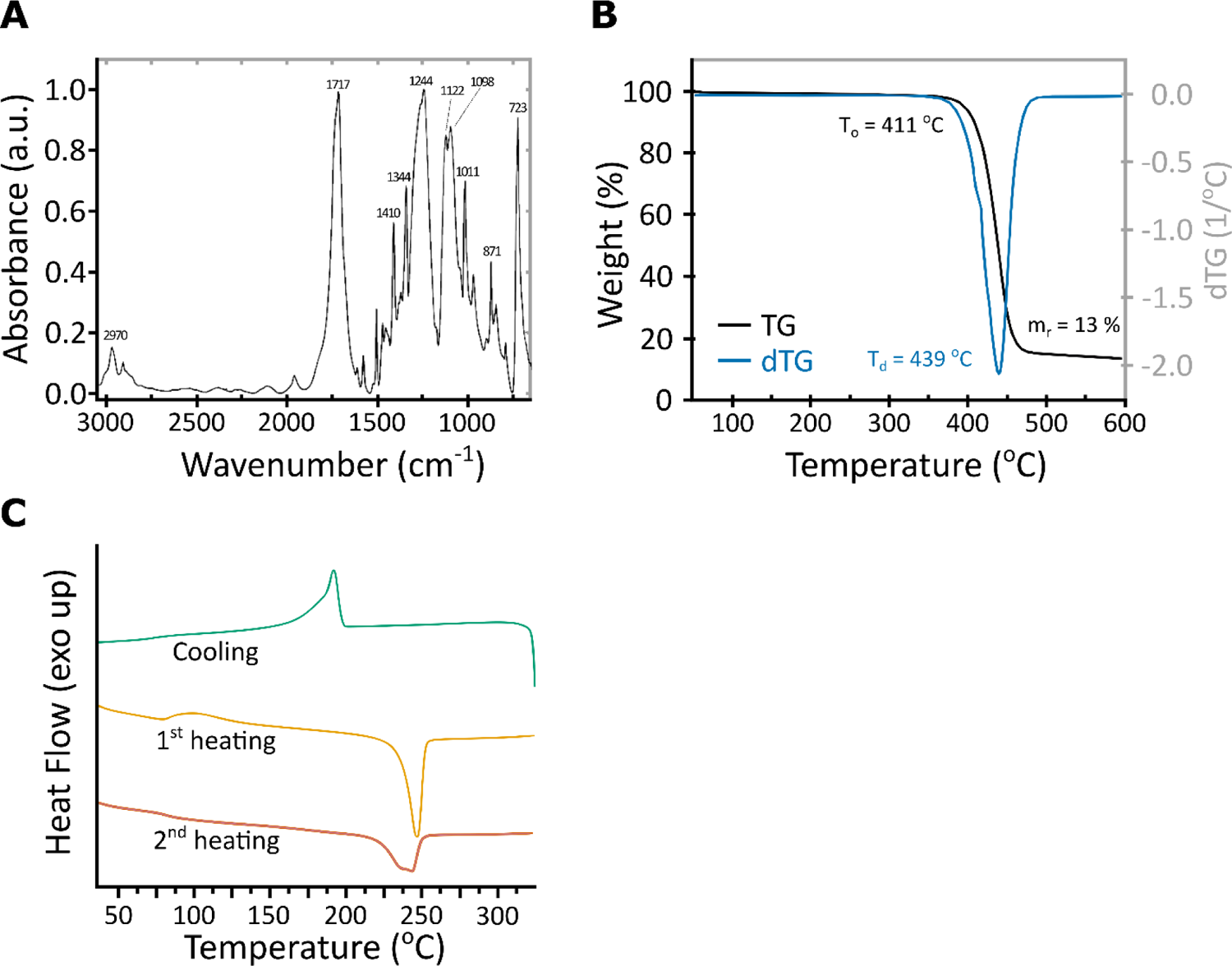
Characterisation of PET plastic: FTIR spectrum of the fabricated PET particles with marked wavenumbers of the peaks of interest (A). TG (black) and dTG (blue) thermograms of PET particles (B). DSC thermogram of PET particles (C) with representation of the cooling (green), 1st heating (yellow) and 2nd heating (orange) stages.

To confirm that, the thermal stability of the sample can be examined. The TGA results show a single step of the sample decomposition, indicating again that no significant amount of additives is detected (Figure 2B)(Han et al., 2018). The degradation started at the onset temperature (T_o_) equal to 411 °C, with the maximum rate at 439 °C, as indicated by the peak of the derivative thermogravimetric (dTG) curve. The residual mass was equal to 13 %, which in the case of the measurement performed in nitrogen atmosphere is attributed to the char.

The DSC results show a typical for PET behavior during the 1^st^ heating with a glass transition at 86 °C and an endothermic peak corresponding to melting at 247 °C (Figure 2C)(Antoniadis et al., 2011). Then, during cooling, an exothermic peak correlated with crystallization is observed at 192 °C. The peak is sharp at the higher temperature, but there is a slight shoulder at 182 °C, indicating the crystallization of a different population, such as chains with a different molecular weight. The glass transition is noted at 76 °C. Finally, during the 2^nd^ heating, the glass transition can be seen at 82 °C, followed by a bimodal melting peak with the maxima, at 237 and 244 °C. This is in agreement with the crystallization peaks, and again, two populations of PET chains, possibly with different molecular weights.

### 3.2. Cell survivability

Cells counting was performed manually using a hemocytometry to assess whether 6 days exposure of PET at 50 ppm induces cell death, as nanoparticles can interfere with cell viability assays (Ong et al., 2014).

There is a reduction of number of pericytes following 6 days treatment of PET at 50 ppm (Unpaired t-test: p = 0.0427, F (5,5) = 13.92, n = 6 per experimental group), representing an overall decrease of 10.2 % cell population compared to controls.

### 3.3. Oxygen consumption rate (OCR) and extracellular acidification (ECAR)

The Agilent Seahorse XFp Mito Stress Test was used to assess different bioenergetic measures of the mitochondrial respiration. The Mito Stress Test measures the OCR expressed in pmol/min and ECAR expressed in mpH/min. There is a n = 5 and n = 6 for the control and the PET experimental group respectively.

ECAR expressed in mpH/min (2-way ANOVA; Experimental group: p = 0.0027, F (1,9) = 16.68; Time: p < 0.0001, F (2.863, 25.76) = 1094; Well: p < 0.0001, F (9,99) = 171.9; Interaction: P < 0.0001, F (11,99) = 6.685; Figure 3A) and the OCR expressed in pmol/min ((2-way ANOVA; Experimental group: p = 0.0010, F (1,9) = 23.08; Time: p < 0.0001, F (1.340, 12.06) = 879.7; Well: p < 0.0001, F (9,99) = 8.432; Interaction: p < 0.0001, F (11,99) = 17.71; Figure 3B) were both increased in pericytes exposed to 6 days of PET compared to the control group.

**Figure 3.**
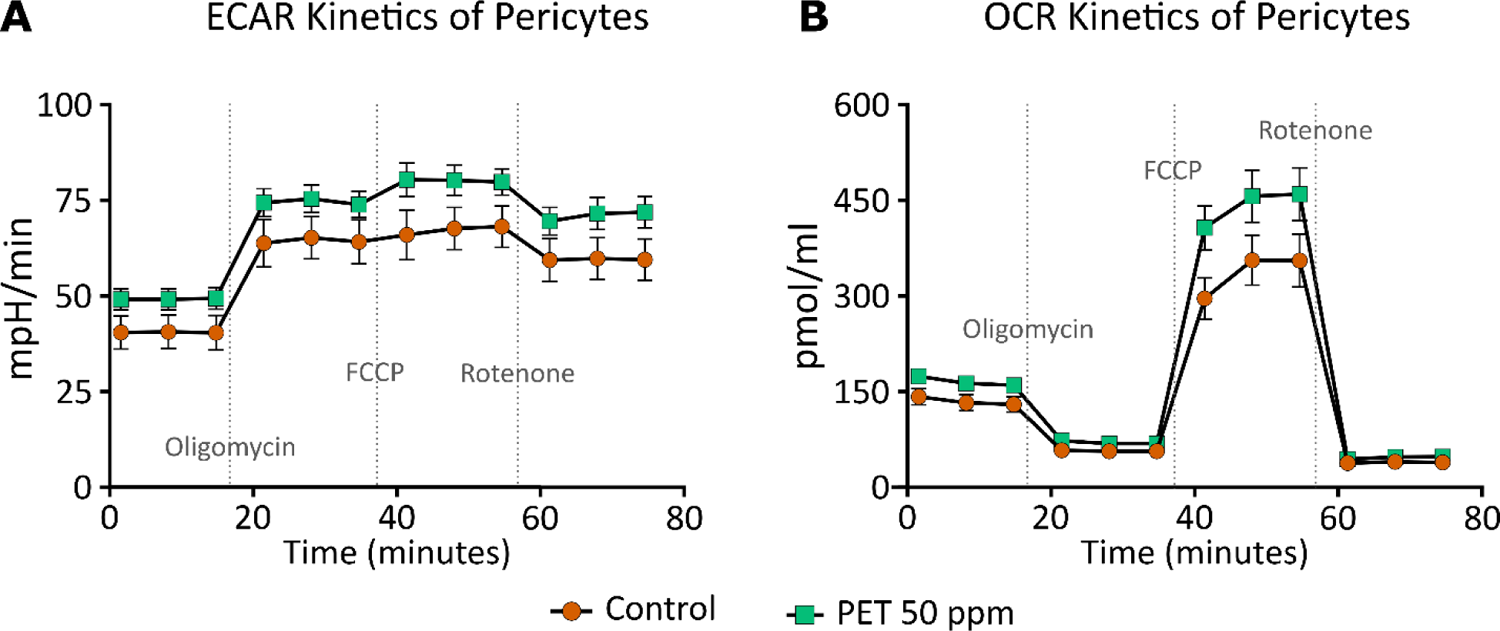
Extracellular acidification rate (ECAR, A) and oxygen consumption rate (OCR, B) kinetics: ECAR and OCR of pericytes exposed to control (round orange) or 50 ppm PET (square green) for 6 days.

The non-mitochondrial oxygen consumption (unpaired t-test: p = 0.0505, F (5,4) = 4.439; Figure 4A), the basal respiration (Mann-Whitney test: p = 0.0823, Mann-Whitney U = 5; Figure 4B), and the coupling efficiency expressed in percentage (unpaired t-test: p = 0.4314, F (5,4) = 2.565; Figure 4C) were not statistically significant between both experimental groups, yet it is worth noting that pericytes exposed to 6 days of PET had a tendency (p = 0.505) to have a higher non-mitochondrial oxygen consumption.

**Figure 4.**
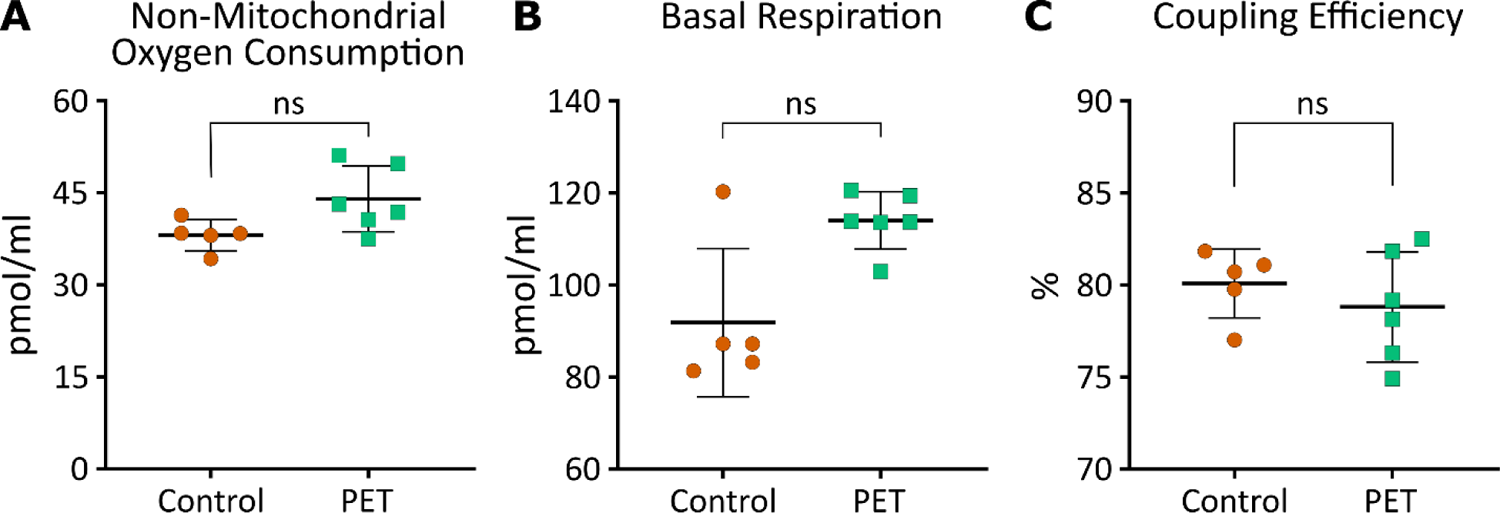
Non-mitochondrial oxygen consumption (A), basal respiration (C) and coupling efficiency (C) activity. Control (round orange) and 50 ppm PET (square green) exposure for 6 days on pericytes. ns = statistically non-significant P > 0.05.

The pericytes exposed to PET for 6 days had a higher proton pump leak (Mann-Whitney test: p=0.0303, Mann-Whitney U = 3; Figure 5A), ATP production (unpaired t-test: p = 0.0040, F (5,4) = 1.106; Figure 5B), maximal respiration (unpaired t-test: p = 0.0031, F (5,4) = 1.076; Figure 5C) and spare respiratory capacity (unpaired t-test: p = 0.0037, F (5,4) = 1.146; Figure 5D) compared to the control group.

**Figure 5.**
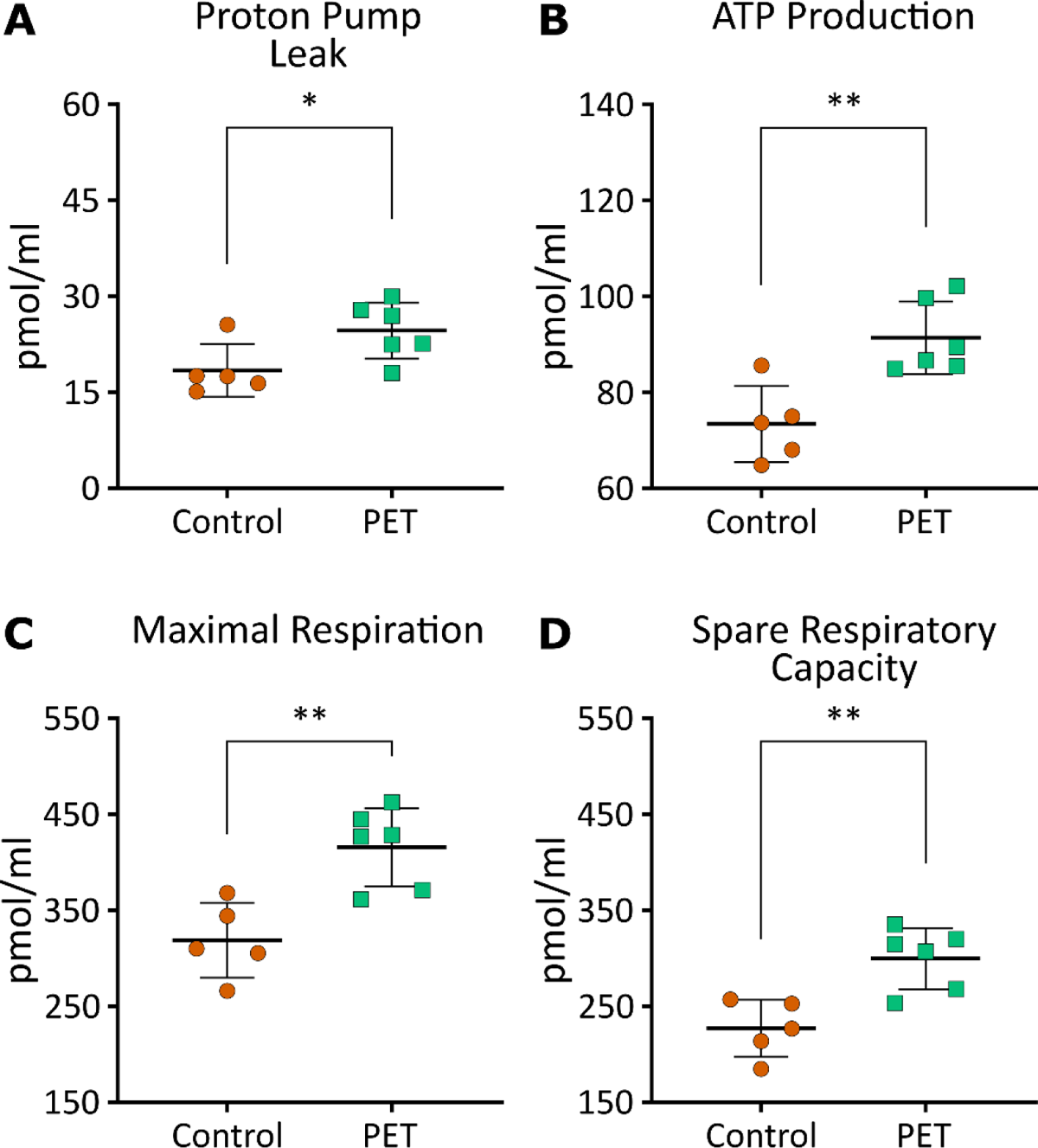
Proton pump leak (A), ATP production (B), maximal respiration (C) and spare respiratory capacity (D) activity. Control (round orange) and 50 ppm PET (square green) exposure for 6 days on pericytes. *P ≤ 0.05; ** P ≤ 0.01.

### 3.4. Mitochondrial copy

To assess whether PET exposure can affect the number of mitochondria of pericytes, relative mitochondrial DNA copy was measured using qPCR. No significant difference was noted in the relative mitochondrial DNA copies between both experimental groups (Mann-Whitney test: p = 0.6857; n = 4 per group).

### 3.5. ROS level

To detect the presence of ROS, the general oxidative stress indicator CM-H2DCFDA was used, and the geometric mean fluorescence intensity was measured. Excitation of the probe was ∼492–495 nm whereas it emitted wavelengths of 517–527 nm. Pericytes exposed to 50ppm PET for 6 days did not show a significant difference in ROS expression compared to control exposure (Unpaired t-test: p = 0.1314, F (4,4) = 2.690, n = 5 per experimental group).

### 3.6. Expression of genes associated with oxidative stress and ferroptosis

The level of expression of genes associated with oxidative stress and ferroptosis were assessed using RT-qPCR. There was no difference in the expression level of genes associated with oxidative stress, ROS or ferroptosis, when comparisons were drawn between the control group and the PET-exposed pericytes (Table 2). While *NCOA4* was statistically increased in pericytes exposed to PET particles at 50 ppm for 6 days, the change was only of an average of 1.43-fold with a standard deviation of 0.27 compared to the control group at an average of 1 with a standard deviation of 0.20, this suggests that the increase not physiologic as this is within the variation of the RT-qPCR technique. Unpaired t-test was used with an n = 6 for both experimental groups.

**Table 2.** The ΔCT, standard deviation (SD), p, and F values of gene expression (as a function of detected mRNA) levels of pericytes exposed to 50 ppm PET for 6 days compared against untreated controls.

| Genes | Control ΔCT value<br>(mean ± SD) | PET ΔCT value<br>(mean ± SD) | p<br>value | F (5,5) |
| --- | --- | --- | --- | --- |
| *NOTCH3* | 4.86 ± 0.54 | 4.78 ± 0.36 | 0.7601 | 2.259 |
| *SOD1* | 1.40 ± 0.74 | 1.07 ± 0.31 | 0.3386 | 5.646 |
| *HIF1α* | -0.80 ± 0.46 | -1.21 ± 0.25 | 0.0858 | 3.502 |
| *GXP4* | 2.07 ± 0.36 | 1.74 ± 0.38 | 0.1593 | 1.121 |
| *SLC7A11* | 6.60 ± 0.43 | 6.56 ± 0.49 | 0.8783 | 1.284 |
| *NCOA4* | 1.74 ± 0.29 | 1.22 ± 0.28 | 0.0105 | 1.047 |
| *KEAP1* | 3.69 ± 0.24 | 3.29 ± 0.59 | 0.1592 | 6.033 |

## 4. Discussion

The aim of this study was to assess whether PET particles at 50 ppm would have an impact on monocultured pericyte cells *in vitro*. We found that pericytes do not show sign of oxidative stress as indicated by no ROS changes nor changes in expression level of gene associated with oxidative stress as a result of PET nanoparticle exposure at this level. This is unexpected as cells cultured in lower concentrations of FBS, and cells that have the capability to phagocytose are more susceptible to cytotoxicity from plastic particles (Fröhlich et al., 2012). These pericytes were cultured with 2 % FBS also had the ability to phagocytose (Winkler et al., 2014).

The targeted genes where selected for their implication in the oxidative stress pathways and ferroptosis (Li et al., 2020), which is the iron-dependant cell death connected to ROS signalling. Amongst protective signalling pathway that reduces ROS level, oxidative stress or ferroptosis, we have selected *NOTCH3* for protection from increase of ROS and ferroptosis (Li et al., 2022), *HIF-1*α (Li et al., 2019) and *SOD1* (Eleutherio et al., 2021) for their ability to reduce ROS levels and oxidative stress. Amongst genes involved in the ferroptosis pathway we selected *GXP4* and *SLC7A11* both being anti-ferroptosis biomarkers (Zuo et al., 2020) part of the ferroptosis GSH/GPX4 pathway, and *NCOA4* and *KEAP1* both associated to the iron metabolism (Li et al., 2020) and oxidative stress (Del Rey and Mancias, 2019; Yu and Xiao, 2021).

Additionally, the age of the polystyrene plastic particles influences their toxicity (Murali et al., 2015), as “aged” polystyrene nanoplastics aggregates more compared to fresh polystyrene nanoplastics and also accumulated adsorption of bioactive compounds. In this experiment, freshly made PET particles were produced and utilised rather than “aged” PET particles (> 6 months) which could explain the lack oxidative stress effects.

ECAR, OCR and some bioenergetics aspects of the mitochondrial functions are increased by PET particles at 50 ppm exposure for 6 days. ECAR measures the total acidity produced from glycolysis and the tricarboxylic acid cycle, therefore it is not possible to distinguish the exact source of the acidity. Extracellular acidification has been shown to induce ROS, leading to cell death (Teixeira et al., 2018), however neither oxidative stress, nor changes in mitochondrial copies was noted, despite a 10 % cell death, but cell counting was done manually using a hemacytometer with a statistical decrease of cell population (p = 0.042) near the p < 0.05 threshold, therefore we suggest applying cautiousness on whether PET particles at 50 ppm is cytotoxic per say. It could be hypothesised that the cell death observed is a statistical event that has no physiological relevance to the variables in this experiment.

Interestingly, an increase of mitochondrial functions induced by 6 days exposure of PET particles at 50 ppm was observed, therefore the absence of oxidative stress might not be caused by lack of PET effect but rather mitohormesis (Yun and Finkel, 2014) induced by PET.

Mitohormesis refers to the protective mechanism of which a mild mitochondrial stressor induces a mitochondrial adaptive response (hormetic response) to then protect the cell against future stressors. The pericyte bioenergetic changes induced by PET exposure show that there is a higher demand of ATP with a higher ATP production and a higher respiratory capacity suggesting a physiological adaptation (Hill et al., 2012) which may be mitohormesis in response to the stress induced by PET exposure. Despite ROS signalling being a part of the mitohormesis response (Bárcena et al., 2018), we did not observe increases of ROS after 6 days exposure of PET, which is not surprising as ROS signalling involves feedback loop with eventually a reduction of ROS while the increased mitochondrial activity is maintained (Zarse et al., 2012).

Micro- and nanoplastics would not be the first toxic substance to induce mitohermesis; low-dose arsenite has been show to induce mitohormesis thus increasing lifespan (Schmeisser et al., 2013) and could potentially be linked to reduced risk of cancer (Ahn et al., 2020; Baastrup et al., 2008). However, it is important to understand at which dose PET might provide a beneficial effect via hormesis. This is because mitohormesis can be represented as a U-shaped curve where very low-doses and very high-doses of ROS are harmful but low-doses of ROS may be beneficial (Kim et al., 2018; Lee and Lee, 2019). This experiment only probed into one dose of PET at 50 ppm, while an investigation into plastic particles in human blood suggested that average plastic contamination could be as low as 1.62 (Leslie et al. 2022), not accounting for occupational exposure and bioaccumulation. Leslie et al. 2022 examined 22 anonymised adult donors and therefore we hypothesise that this might not reflect individuals subject to higher plastic exposures, such as occupational exposures. For instance, 3D printing has been an increasing source of concerned related to indoor air pollution (Salthammer, 2022). Additionally, bioavailability and accumulation of nanoplastics in the brain is unknown, yet accumulation of particle matters of diameter less than 1 µm could be the cause of BBB damage (Shih et al., 2018). Therefore, we have used a higher concentration of plastic particles to consider higher exposures and potential accumulation at the BBB.

An interesting derivative experiment looking into lower doses of PET may reveal more findings regarding mitohormesis. Further studies are needed to establish potential accumulations of nanoplastics in the blood brain barrier as well as the effects on the surrounding milieu to ascertain a physiologically relevant chronic dose and to address this toxicologic hormetic dose-response relationship (Calabrese, 2005).

The choice of plastics and concentrations must be evaluated and considered. Ready-to-use nano and microplastics suspensions are smooth rather than being rough surfaced which does not correspond to realistic human exposure (de Sá et al., 2018), and they are often sold mixed with additive agents that can induce toxic activity on their own. Similarly, the high concentration of plastic particles used in *in vitro* studies can interfere with the cell medium and toxicity (Stock et al., 2022). This raises the question of whether plastic particles or organic additives to the plastics are more of a health concern. For instance, it has been recently shown that the additive compounds used with plastics can be more cytotoxic than the microplastic particles; however, this is dependent on size and surface charge. Stock and colleagues (2022) demonstrated that the cytotoxicity and cellular uptake of nanoplastics correlated directly with their size and surface charge, where smaller nanoparticles were taken up quicker and were more cytotoxic. In our study, we confirmed that we obtained plastics of PET by FTIR spectroscopy. No significant amount of additives was observed, as confirmed also by TGA studies. The PET particles were most likely composed of two different populations with varying molecular weights, as shown by DSC experiments. The formation of two populations of PET could be explained by the methods used to make plastic particles from PET jars, which included grinding, milling and sieving techniques under different temperature ranges (room temperature, dry ice, liquid nitrogen).

This *in vitro* study aimed at focusing on the effect of plastic particles directly on pericyte cell only. The impact of nanoplastic particles on the blood-brain-barrier, nor the interplay of pericytes with other cell types to carry out their complex functions, was investigated. For example, impaired pericytes reduce the blood-brain barrier function and induce toxin accumulation in the brain (Bell et al., 2010). Therefore, more studies are needed to understand the effects of nanoplastic particles on the complex blood-brain barrier using a mouse organotypic cortical culture (Bendfeldt et al., 2007) or a human induced pluripotent cells (Mesentier-Louro et al., 2023) model of the blood-brain barrier, and whether PET exposure would induce similar effects in other cells involved in a healthy blood-brain-barrier.

## 5. Conclusion

This study demonstrates that *in vitro* exposure of a monoculture of human pericytes to 50 ppm PET particles for 6 days induced 10 % cell death and potential mitohormesis. Critically, whilst oxidative stress was not indicated, increased proton pump leak, maximal respiration, spare respiratory capacity, ATP-production, and ECAR/OCR were noted. Further research looking into the stages of mitohormesis that may be being induced is necessary to definitively claim that PET particles may induce mitohormesis.

Derivative experiments looking at the integrity of models of the blood-brain-barrier, as well as intercellular interactions between pericytes and other cells forming the BBB, for example endothelial cells and astrocytes, are necessary to shed light on the full capabilities of plastic particles to induce potentially toxic and impairing effects or mitohormesis. Research into the accumulation of nanoplastics in the milieu of the blood-brain-barrier would shed light on the impact of nanoparticles on the tight junctions of the endothelial cells, for example.

In summary, although further research is needed, this investigation implicates PET particles in impacting mitochondrial function in human brain vascular pericytes.

### CRediT authorship contribution statement

**Sean M. Gettings**: Investigation, Methodology, Resources, Visualisation, Validation, Writing - review and editing, Conceptualisation, Formal analysis, Supervision, Funding acquisition, Project administration. **Will Timbury**: Investigation, Methodology, Resources, Visualisation, Validation, Writing - review and editing. **Anna Dmochowska**: Investigation, Methodology, Visualisation, Validation, Writing - review and editing, Conceptualisation, Formal analysis. **Riddhi Sharma**: Investigation, Methodology, Resources, Visualisation, Validation, Writing - review and editing. **Lewis E. MacKenzie**: Investigation, Methodology, Resources, Visualisation, Validation, Writing - review and editing, Conceptualisation, Formal analysis. **Guillaume Miquelard-Garnier**: Methodology, Resources, Writing - review and editing, Conceptualisation, Supervision. **Nora Bourbia**: Investigation, Methodology, Resources, Visualisation, Validation, Writing – original draft, Conceptualisation, Formal analysis, Supervision, Funding acquisition, Project administration.

## Acknowledgement

This study is part funded by the National Institute for Health and Care Research (NIHR) Health Protection Research Unit in Chemical and Radiation Threats and Hazards (NIHR 200922), a partnership between UK Health Security Agency and Imperial College London. The views expressed are those of the author(s) and not necessarily those of the NIHR, UK Health Security Agency or the Department of Health and Social Care.

Lewis MacKenzie was supported by a BBSRC Discovery Fellowship award (BB/T009268/1). The Ecole Doctorale SMI (ED 432) is acknowledged for granting Anna Dmochowska the fellowship for her Ph.D. work. We thank Lorna Jones (UKHSA) for her advice in the making of plastic particles. We thank Dr Paul R. Edwards and Professor Robert W. Martin (Department of Physics, University of Strathclyde) for access to and support with STEM and SEM imaging.

